# Opioid withdrawal abruptly disrupts amygdala circuit function by reducing peptide actions

**DOI:** 10.1101/2021.12.22.471860

**Authors:** Gabrielle C. Gregoriou, Sahil D. Patel, Sebastian Pyne, Bryony L. Winters, Elena E. Bagley

## Abstract

Opioid withdrawal drives relapse and contributes to compulsive drug use through disruption of endogenous opioid dependent learning circuits in the amygdala. Normally, endogenous opioids control these circuits by inhibiting glutamate release from basolateral amygdala principal neurons onto GABAergic intercalated cells. Using patch-clamp electrophysiology in rat brain slices, we reveal that opioid withdrawal dials down this endogenous opioid inhibition of synaptic transmission. Peptide activity is dialled down due to a protein kinase A dependent increase in the activity of the peptidase, neprilysin. This disrupts peptidergic control of both GABAergic and glutamatergic transmission through multiple amygdala circuits, including reward-related outputs to the nucleus accumbens. This likely disrupts peptide-dependent learning processes in the amygdala during withdrawal. and may direct behaviour towards compulsive drug use. Restoration of endogenous peptide activity during withdrawal may be a viable option to normalise synaptic transmission in the amygdala and restore normal reward learning.

**In Brief:** We find that opioid withdrawal dials down inhibitory neuropeptide activity in the amygdala. This disrupts both GABAergic and glutamatergic transmission through amygdala circuits, including reward-related outputs to the nucleus accumbens. This likely disrupts peptide-dependent learning processes in the amygdala during withdrawal and may direct behaviour towards compulsive drug use.

**Highlights:**

- During opioid withdrawal, peptidase activity is upregulated in an amygdala circuit
- Peptidase upregulation occurs via a PKA-dependent mechanism
- Increased peptidase activity limits peptidergic control of synaptic transmission
- Opioid withdrawal disrupts the balance of excitation and inhibition in the amygdala

## Introduction

Opioid addiction is a complex brain disorder, characterised by recurring cycles of compulsive drug-taking, withdrawal, and relapse, even after prolonged periods of abstinence (Koob and Volkow, 2016). Long-term propensity to relapse is associated with persistent changes in reward circuits following chronic drug use, such that the systems that shape adaptive behaviours become dysregulated and progressively directed toward compulsive drug-seeking and drug-taking (Lüscher et al., 2020). The basolateral amygdala (BLA) is a critical node within the brain’s reward circuits (Murray, 2007, Janak and Tye, 2015). Multimodal information related to the pursuit of rewards and/or the stimuli that predict them converge on specific BLA principal neurons, which link these discrete actions and/or stimuli to the reward’s positive affective outcome (Namburi et al., 2015, Paton et al., 2006). BLA principal neurons encode lasting mental representations of these learned outcomes and can retrieve them when necessary to guide adaptive behaviour (Wassum and Izquierdo, 2015) through their projections to other brain regions, such as the nucleus accumbens (NAc). Outcome representations are dynamic and can be updated with respect to state-dependent changes in reward incentive value (Belova et al., 2008, Wassum et al., 2009, Salzman et al., 2007), which is also encoded and retrieved by BLA principal neurons (Malvaez et al., 2019). For example, exposure to a reward, such as opioid or food, during opioid withdrawal elevates the value of the reward experienced during this time (Wassum et al., 2016, Hutcheson et al., 2001). This elevated value persists beyond the period of withdrawal and increases subsequent responding to any associated cues (Wassum et al., 2016, Hutcheson et al., 2001), which may facilitate the development of compulsive drug use. This suggests that opioid withdrawal may open a window where disrupted learning in BLA circuits occurs. This window may be opened by adaptations that develop in opioid-sensitive neurons during chronic opioid use and are unmasked when drug use is ceased (Williams et al., 2001). As reward re-evaluation relies on endogenous opioids acting at the µ-opioid receptor in the BLA (Wassum et al., 2016), adaptations within the endogenous opioid system will likely influence this learning.

The endogenous opioids that regulate reward learning in the BLA during opioid withdrawal are likely sourced from neighbouring GABAergic intercalated cell (ITC) clusters. A main island (Im) of these cells, which sit at the base of the BLA and are activated by BLA principal neurons, provide direct inhibition back onto BLA principal neurons. Through these functional connections, Im neurons are able to regulate activity of BLA principal neuron outputs (Asede et al., 2015), and ultimately BLA principal neuron-mediated learning and memory processes (Busti et al., 2011, Jüngling et al., 2008, Likhtik et al., 2008). We have recently shown that Im neurons release endogenous opioids that inhibit neurotransmitter release from both BLA principal neurons and Im neurons, thereby directly inhibiting neuronal excitability within the Im (Winters et al., 2017). A major controller of endogenous opioid function in the amygdala is the peptidases that break down opioid peptides into inactive fragments, with inhibition of these enzymes doubling endogenous opioid actions (Winters et al., 2017). In several brain regions, including the periaqueductal gray and striatum, biochemical measures of enkephalin-degrading peptidase activity are increased during opioid withdrawal (Malfroy et al., 1978, Zhou et al., 2001). It is possible that these widespread increases in peptidase activity represent an adaptation to chronic opioids that is common throughout opioid-sensitive circuits. Im ITCs are likely to develop adaptations to chronic opioid treatment, such as increases in peptidase activity, because they express very high levels of both the µ-opioid receptor (Poulin et al., 2006). If increased peptidase activity is unmasked in the amygdala during opioid withdrawal, peptidergic control of synaptic transmission and cellular excitability during this period would be reduced. This would disrupt the balance of ITC-BLA circuits and dysregulate the BLA principal neuron-mediated learning processes that drive maladaptive drug-seeking and relapse behaviours during withdrawal and abstinence.

In this study, we aimed to determine whether opioid withdrawal alters peptide regulation of amygdala synapses. Whole-cell patch recordings of BLA principal neurons and Im ITCs from brain slices spontaneously withdrawn from morphine were used. We demonstrate that opioid withdrawal reduces peptide control of synaptic transmission between BLA principal neurons and ITCs. This reduced peptidergic control is dependent on protein kinase A (PKA)-driven upregulation of peptidase activity and alters control of reciprocal BLA-Im synapses, including BLA neurons that project to the NAc. Thus, we establish that amygdala-mediated processes are vulnerable during withdrawal, and suggest that withdrawal opens a window whereby the long-term modifications in reward learning that rely on endogenous opioid actions (Wassum et al., 2016) and contribute to the development of compulsive drug use can be laid down.

## Results

### Opioid withdrawal reduces MOR, but not DOR, inhibition of glutamate release at BLA-Im ITC synapses

To induce morphine dependence, Sprague Dawley rats were chronically treated with a slow-release morphine emulsion (Bagley et al., 2011) and control rats were treated with a morphine-free vehicle emulsion on the same schedule (Figure 1A). Brain slices taken from these animals were incubated in either a bath of artificial ACSF containing morphine (5µM) to study the effects of chronic morphine treatment or a morphine-free bath to study the effects of spontaneous *in vitro* morphine withdrawal (Bagley et al., 2005) (Figure 1A). To test whether this chronic morphine treatment induced opioid dependence we examined whether injection of the opioid receptor antagonist naloxone (5mg/kg) increased opioid withdrawal signs (Figure 1B). Neither vehicle-treated control animals nor morphine dependent animals displayed any signs of withdrawal prior to the naloxone challenge (Figure 1C) and no withdrawal signs were observed in vehicle-treated rats after the naloxone challenge (Figure 1C). In contrast, after chronic morphine-treatment rats exhibited all the counted withdrawal signs post-naloxone injection (Figure 1C). This indicates that, as seen previously (Bagley et al., 2011, Bagley et al., 2005), the current chronic morphine treatment protocol successfully induced opioid dependence.

**Figure 1.**
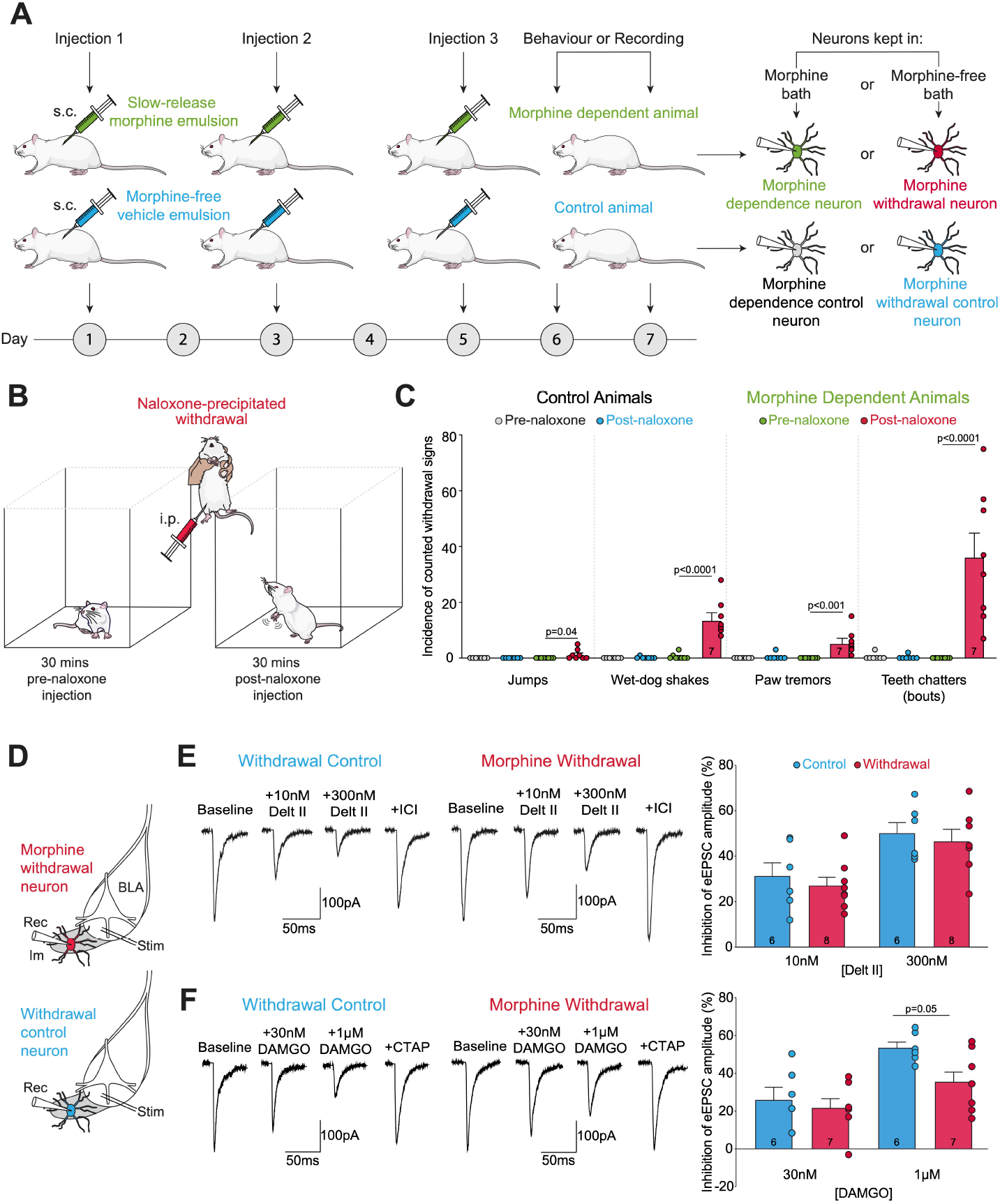
Opioid withdrawal reduces MOR, but not DOR, inhibition of glutamate release at BLA-Im ITC synapses. A) Schematic of the dependence paradigm showing that a slow-release emulsion containing 50mg/kg morphine was injected into rats every second day to induce dependence. Control rats were injected with a morphine-free vehicle emulsion on the same schedule. Morphine dependent and control animals were used for experiments on days 6 or 7. Brain slices taken from these rats were incubated in either a bath containing 5µM morphine or a morphine-free bath to study morphine dependence or spontaneous *in vitro* withdrawal, respectively. B) Schematic showing that rats were observed for 30 mins pre- and post-injection of 5mg/kg naloxone hydrochloride. Withdrawal signs including, jumps, wet-dog shakes, paw tremor, and teeth chattering were counted during each observation period. C) In morphine dependent animals (n=8), naloxone injection significantly increased all withdrawal signs. Control animals (n=7) displayed no signs of withdrawal neither pre-nor post-naloxone injection. D) Representative BLA *stim*ulation and Im *rec*ording locations. E) The selective DOR agonist Delt II reduced eEPSC amplitude to a similar extent in control and morphine withdrawn neurons at both submaximal and maximal concentrations. Example eEPSCs of single representative experiments. Bar chart showing percentage inhibition from baseline. F) The selective MOR agonist DAMGO reduced eEPSC amplitude in both control and morphine withdrawn neurons, however, there was a significant effect of treatment group on percent inhibition by DAMGO (F_1,23_=4.32, p=0.05, 2-way ANOVA), with the effect of DAMGO at maximal concentration reduced in morphine withdrawn neurons compared to control. Example eEPSCs of single representative experiments. Bar chart showing percent inhibition from baseline. Bar charts show mean ± SEM. Circles represent individual neurons. Neuron number is shown in bars. Data were analysed using 2-way ANOVAs with comparisons on graphs showing results from post-hoc Bonferroni’s multiple comparisons tests. See Table S1 for full statistics and analysis.

To study whether opioid withdrawal alters peptidase control of neuropeptide activity in the amygdala our initial experiments focused on regulation of the BLA principal neuron synapse onto Im neurons (BLA-Im synapse). We chose to study this synapse as peptidases strongly limit inhibition of this synapse by both exogenous and endogenous enkephalins in untreated rats (Winters et al., 2017, Gregoriou et al., 2020). Our measurement of peptidase activity during withdrawal relied upon unchanged opioid receptor signalling, however, in other brain regions, withdrawal can either increase or decrease opioid receptor coupling to their effectors (Ingram et al., 1998, Ingram et al., 2008, Hack and Christie, 2003, Bagley et al., 2005, Chieng and Christie, 1996). To control for the possibility of withdrawal-induced changes in opioid receptor signalling, we initially tested whether spontaneous withdrawal from chronic morphine treatment induced changes to opioid receptor coupling to inhibition of glutamate release at BLA-Im synapses. To do this we electrically stimulated the BLA (Figure 1D) and recorded the resulting evoked excitatory postsynaptic currents (eEPSC) in Im neurons during whole cell patch clamp recordings. As activation of μ- and δ-opioid receptors (MORs and DORs), but not κ-opioid receptors (KORs), inhibits glutamate release at BLA-Im synapses (Winters et al., 2017), selective agonists for MORs and DORs were used to test opioid receptor coupling at this synapse. Non-hydrolysable MOR and DOR agonists were chosen so that drug effects were independent of peptidase activity. Deltorphin II (Delt II), a selective and non-hydrolysable agonist at the DOR, was tested at a submaximal (10nM) and maximal concentration (300nM) (Hack et al., 2005). Delt II inhibited eEPSCs at the BLA-Im synapse in a concentration-dependent manner in both control and morphine-withdrawn neurons (Figure 1E). Morphine-withdrawal did not alter the efficacy of Delt II to inhibit glutamate release at either concentration (Figure 1E). This indicates that withdrawal does not modify DOR coupling at this synapse. The DOR-selective antagonist, ICI-174864 (1μM), reversed the Delt II-induced inhibition in all neurons (Control neurons reversal = 101.91% ± 5.43% (n = 6), Withdrawn neurons reversal = 89.30% ± 4.31% (n = 8)). DAMGO, a selective and non-hydrolysable agonist at the MOR, was also tested at submaximal (30nM) and maximal concentrations (1μM) (Ingram et al., 1998). While submaximal DAMGO inhibited glutamate release to the same extent in both groups, at the maximal concentration, there was a reduction in the effect of DAMGO in morphine-withdrawn neurons (Figure 1F). The MOR antagonist CTAP (1μM) reversed the DAMGO inhibition of glutamate release in all cells (Control neurons reversal = 97.25% ± 3.96% (n = 6), Withdrawn neurons reversal = 100.01% ± 3.28% (n = 8)). These data indicate that spontaneous withdrawal from chronic morphine treatment reduces the ability of MORs to regulate glutamate transmission from BLA principal neurons to Im ITCs while DOR-dependent regulation of glutamate release remains unaffected. This suggests MOR activity on BLA terminals is reduced after chronic treatment and withdrawal from the MOR-preferring agonist, morphine. To prevent this change from distorting our interpretation of peptidase function, all subsequent experiments (unless otherwise stated) were conducted in the presence of the MOR antagonist CTAP (1μM) and thus only opioid actions at DOR on BLA synaptic terminals were measured.

### Neprilysin activity at the BLA-Im synapse is upregulated during opioid withdrawal

The ability of endogenously released enkephalins to inhibit Im neurons and their associated synapses (Winters et al., 2017) is limited by the activity of neprilysin (Gregoriou et al., 2020), an enkephalin-degrading peptidase that cleaves the glycine-phenylalanine bond of enkephalin molecules (Hiranuma et al., 1997, Hiranuma and Oka, 1986, Erdös and Skidgel, 1988). In areas with high expression of opioid receptors, such as the striatum and periaqueductal gray, neprilysin is abundantly coexpressed (Waksman et al., 1986). Given that withdrawal from chronic morphine treatment increases biochemical activity of a neprilysin-like peptidase in these same brain regions (Malfroy et al., 1978, Zhou et al., 2001), we hypothesised that elevated neprilysin activity during opioid withdrawal may be a widespread phenomenon, common in cells with overlapping neprilysin and opioid receptor expression. Considering this, we examined whether neprilysin expression overlapped with opioid receptor expression in the Im. We found high immunoreactivity for neprilysin throughout all amygdala nuclei, including within the Im (Figure 2A). We also found high immunoreactivity for MOR in the Im (Poulin et al., 2008, Winters et al., 2017) and moderate immunoreactivity for enkephalins in both the Im and BLA, as previously observed (Poulin et al., 2008, Winters et al., 2017) (Figure 2A). Within the Im, neprilysin co-localised with enkephalins in MOR-expressing neurons (Figure 2A), suggesting that when Im neurons release enkephalins, neprilysin is well positioned to cleave this enkephalin near its release site, thus critically regulating the extent, timing and spread of endogenous enkephalin activity within the Im.

**Figure 2.**
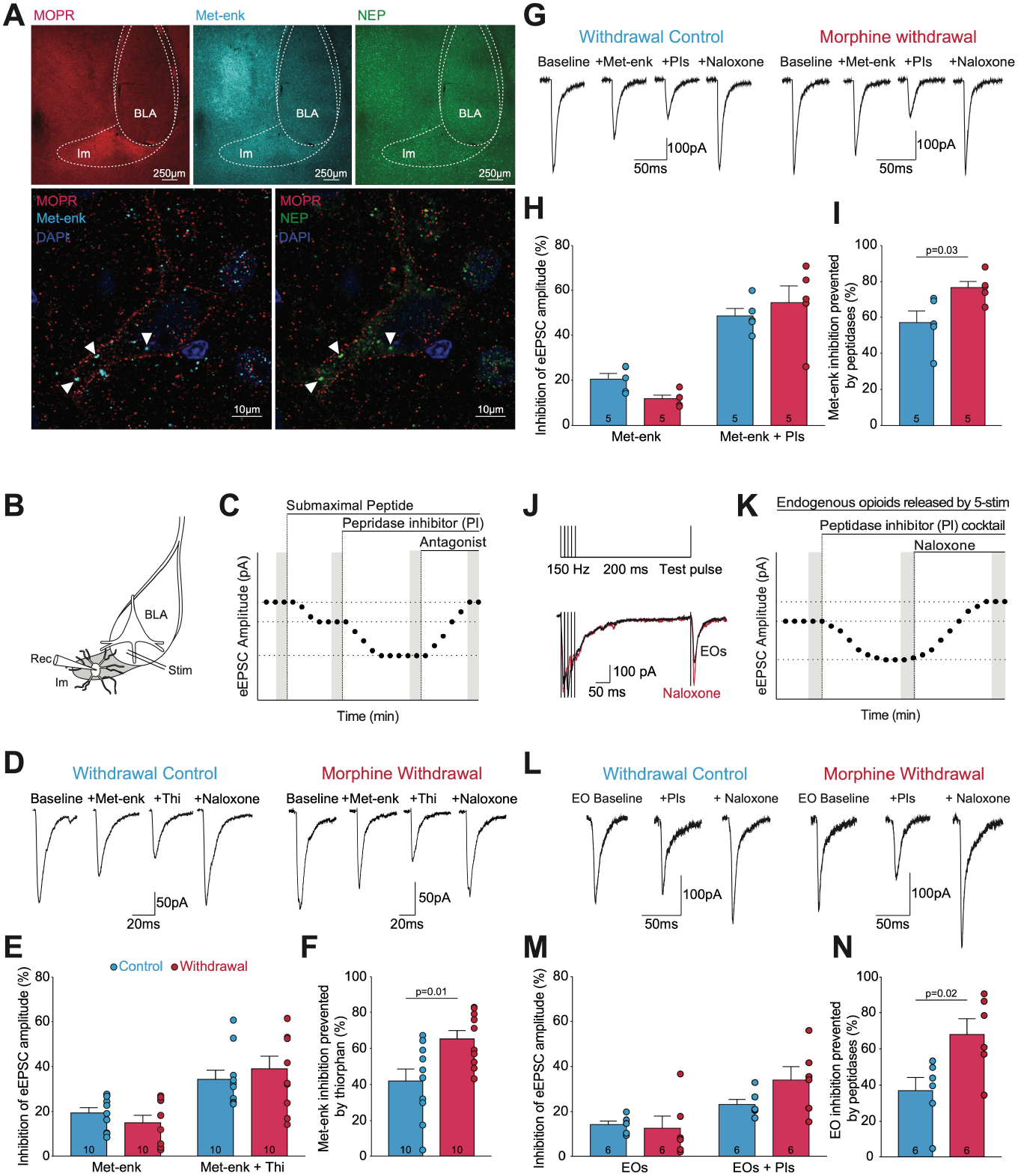
Neprilysin activity at the BLA-Im synapse is upregulated during opioid withdrawal. A) Neprilysin is co-localised with met-enk and MOR in the Im. Top: Low power confocal images showing immunoreactivity for MOR (red), met-enk (blue) and neprilysin (green) throughout the amygdala. Bottom: Magnification of an Im neuron reveals that MOR, met-enk, and neprilysin immunoreactivity co-localises within a single Im neuron. All images were taken at Bregma – 2.00. B-F) Neprilysin regulation of submaximal met-enk (100nM) is enhanced during morphine withdrawal. B) Representative BLA *stim*ulation and Im *rec*ording locations. C) Graphic showing drug application and timing of drug effect measurements (gray bars). D) Example eEPSCs from single representative experiments. E) Bar chart showing percent inhibition from baseline. The neprilysin inhibitor thiorphan (Thi, 10μM) boosted the ability of submaximal met-enk to inhibit eEPSCs in both control and withdrawn neurons (F_1,18_=87.68, p<0.0001, 2-way ANOVA). F) Bar chart showing the proportion of met-enk inhibition prevented by neprilysin activity in control vs. withdrawal. Neprilysin activity was enhanced in opioid withdrawal. G-I) Morphine withdrawal enhanced the ability of a PI cocktail (PIs = thiorphan (10μM), captopril (1μM), bestatin (10μM)) to control submaximal met-enk inhibition. G) Example eEPSCs from single representative experiments. H) Bar chart showing percent inhibition from baseline. The addition of PIs boosted the ability of submaximal met-enk to inhibit eEPSCs in both control and withdrawn neurons (F_1,8_=71.29, p<0.0001, 2-way ANOVA). I) Bar chart showing the proportion of met-enk inhibition prevented by the activity of PI cocktail-sensitive peptidases. The activity of peptidases targeted by our cocktail was greater during opioid withdrawal. J-N) Withdrawal enhanced peptidase control of endogenously released enkephalin actions. J) Stimulus paradigm for endogenous opioid (EO) release using a protocol of a 5-stimuli train followed by a single test pulse. Example traces show eEPSCs evoked by the stimulus paradigm with and without naloxone (10µM). K) Graphic showing drug application and timing of drug effect measurements (gray bars). L) Example test pulse eEPSCs from single representative experiments. M) Bar chart showing percent inhibition of test pulse from baseline. The PI cocktail boosted the ability of endogenously released opioids to inhibit eEPSCs in both control and withdrawn neurons (F_1,10_=73.00, p<0.0001, 2-way ANOVA). N) Bar chart showing the proportion of endogenous opioid inhibition prevented by cocktail-sensitive peptidases. Peptidase activity was increased during opioid withdrawal. Bar charts show mean ± SEM. Circles represent individual neurons. Neuron number is shown in bars. Data were analysed using 2-way ANOVAs with Bonferroni’s multiple comparisons and unpaired Student’s t-tests. Comparisons on graphs show results from t-tests. See Table S1 for full statistics and analysis.

Opioid withdrawal increases biochemical measures of neprilysin-like activity (Malfroy et al., 1978, Zhou et al., 2001) but whether this alters peptide actions is unknown. If opioid withdrawal increases neprilysin activity in the Im it would likely reduce peptidergic control of synaptic transmission and cellular excitability. To study this, we examined whether inhibition of the BLA-Im synapse by a submaximal concentration of enkephalin (100nM) is limited to a greater degree by neprilysin activity during opioid withdrawal (Figure 2B-F). In both control and morphine-withdrawn neurons, submaximal met-enk inhibited eEPSC amplitude and this inhibition was increased by inhibiting neprilysin with thiorphan (Figure 2D-E). However, in morphine-withdrawn neurons, a significantly larger proportion of the met-enk response was prevented by neprilysin activity (Figure 2F). This suggests that the ability of neprilysin to protect met-enk from degradation is enhanced during opioid withdrawal.

Under normal physiological conditions neprilysin selectively degrades enkephalins in the Im (Gregoriou et al., 2020). However, several other enkephalin-degrading peptidases, such as angiotensin-converting enzyme (ACE) and aminopeptidase N (APN), are expressed in the amygdala (Pollard et al., 1989, Krizanova et al., 2001, Banegas et al., 2005). To examine whether opioid withdrawal alters ACE and/or APN hydrolysis of met-enk we applied a peptidase inhibitor (PI) cocktail to amygdala slices in the presence of submaximal met-enk (100nM). The PI cocktail consisted of thiorphan (10μM) to inhibit neprilysin, captopril (1μM) to inhibit ACE, and bestatin (10μM) to inhibit APN (Gregoriou et al., 2020, Winters et al., 2017). As before, in both control and morphine-withdrawn neurons, submaximal met-enk (100nM) inhibited the eEPSC amplitude, this inhibition was increased by the cocktail of PIs, and a greater proportion of the met-enk inhibition was prevented by peptidase activity during opioid withdrawal (Figure 2G-I). Importantly, the total met-enk inhibition produced in the presence of the PI cocktail was no different from the total met-enk inhibition produced in the presence of thiorphan alone, as seen in Figure 2E (Control cells: Total inhibition with cocktail = 48.53% ± 3.33% (n=5) vs. Total inhibition with thiorphan alone = 34.40% ± 4.01% (n=10), p = 0.18; Withdrawn cells: Total inhibition with cocktail = 54.43% ± 7.72% (n=5) vs. Total inhibition with thiorphan alone = 38.98% ± 5.57% (n=10), p = 0.13; Peptidase effect: F_(1,26)_=6.71, p=0.02, Treatment group effect: F_(1,26)_=0.84, p=0.37, Interaction effect: F_(1,26)_=0.01, p=0.90, 2-way ANOVA with Bonferroni’s multiple comparisons). This indicates that during opioid withdrawal, increased peptidase control of opioid activity is solely due to upregulation of neprilysin activity, not changed ACE or APN activity.

To understand how the increased neprilysin activity during opioid withdrawal might limit the actions of endogenously released peptides we stimulated BLA principal neurons at moderate frequency. This releases enkephalins from the Im at levels limited by peptidase activity, but still sufficient to inhibit neurotransmission from BLA principal neurons (Winters et al., 2017). To determine the level of peptidase control we applied the PI cocktail and to determine the level of endogenous opioid inhibition we applied naloxone (10 µM) at a concentration able to antagonise the DORs mediating the endogenous opioid inhibition (Winters et al., 2017), with CTAP 1µM present throughout the experiment. Naloxone increased eEPSC amplitudes above baseline levels in all neurons indicating that the BLA-Im synapse is under tonic endogenous opioid control (Control neurons: Naloxone increase = 16.93% ± 2.13% (n = 6), Withdrawn neurons: Naloxone increase = 17.15% ± 8.65% (n = 6)). Therefore, the eEPSC amplitude following naloxone superfusion reflected the ‘true baseline’ of glutamatergic signalling at the BLA-Im synapse once endogenous opioid inhibition was blocked (Figure 2K). This value was used to determine the proportion of endogenous opioid signalling that was under peptidase control. We found that endogenously released opioids yielded a small inhibition of eEPSCs, which was enhanced by addition of the PI cocktail in morphine withdrawn neurons (Figure 2L-M). The ability of peptidases to prevent endogenous opioid activity was enhanced in withdrawn neurons compared to control (Figure 2N). These data indicate that, as we saw with exogenous enkephalin, morphine withdrawal increases peptidase control over endogenously released enkephalin activity in the Im.

### Neprilysin upregulation occurs through a PKA-dependent mechanism

Having demonstrated that neprilysin control of endogenous peptide actions at the BLA-Im synapse is upregulated during withdrawal, the following experiments addressed how this occurred. As opioid withdrawal results from cellular adaptations induced by chronic opioid exposure (Williams et al., 2001) two possibilities emerged: 1) neprilysin activity is upregulated during morphine dependence to counteract prolonged occupation of opioid receptors or, 2) is upregulated in response to spontaneous, or CTAP-precipitated, withdrawal from chronic morphine *in vitro*. To address this question, we made recordings from Im neurons in slices that had been maintained in morphine (5μM) or spontaneously withdrawn (Figure 3A). In experiments described above that established neprilysin upregulation during the withdrawal state, the MOR antagonist CTAP was present so that homologous tolerance of MORs would not distort our measurement of peptidase activity. However, this was not possible in the present experiment because CTAP would precipitate withdrawal in the slices maintained in morphine. Therefore, we studied whether peptidase control over the activity of another peptide, nociceptin, which is also controlled by neprilysin at this synapse (Gregoriou et al., 2020), is altered by chronic morphine treatment and/or opioid withdrawal. In withdrawn and control neurons, submaximal nociceptin (100nM) caused a small inhibition of eEPSC amplitude, which was potentiated by thiorphan (10µM) in withdrawn cells (Figure 3B-C). Similar to met-enk, a larger proportion of the nociceptin inhibition was prevented by neprilysin in cells withdrawn from chronic morphine (Figure 3D). In contrast, when slices from morphine-dependent rats or control rats were stored in a morphine-containing bath, the extent to which thiorphan was able to boost synaptic inhibition was no different between groups (Figure 3E-G). Together, these data indicate that increased neprilysin activity is revealed during the withdrawal state rather than during chronic morphine treatment. These data also suggest that withdrawal-induced changes in neprilysin activity can limit the activity of non-opioid peptides.

**Figure 3.**
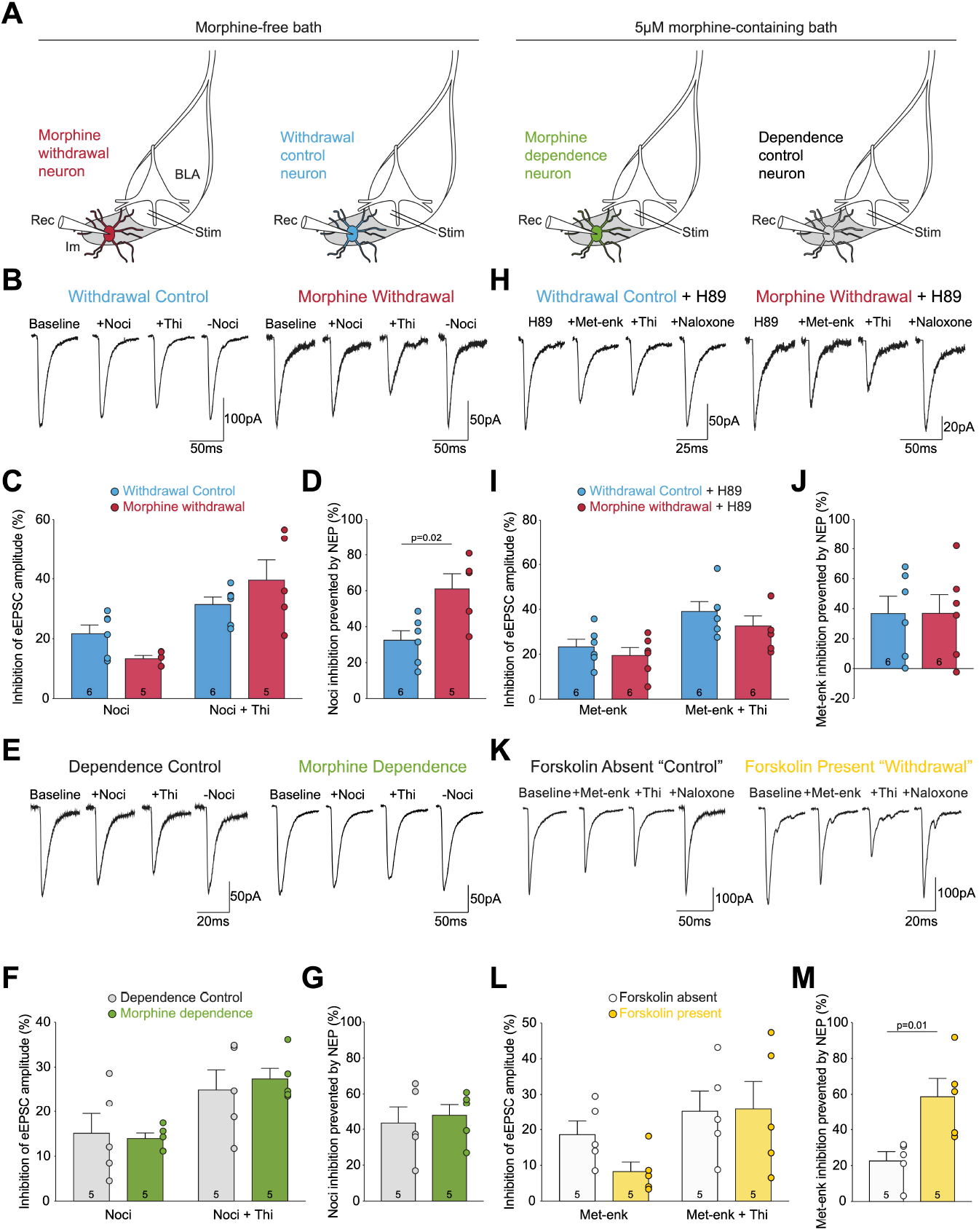
Increased adenylyl cyclase/PKA signalling during opioid withdrawal upregulates neprilysin activity. A) Schematic showing the different experimental groups. B-D) Morphine withdrawal enhances neprilysin control of nociceptin inhibition. B) Example eEPSCs from single representative experiments. C) Bar chart showing percent inhibition from baseline. The neprilysin inhibitor thiorphan (10μM) boosted the ability of submaximal nociceptin (100nM) to inhibit eEPSCs in withdrawn but not control neurons (F_1,9_=29.96, p<0.001, 2-way ANOVA). D) Bar chart showing the proportion of nociception inhibition prevented by neprilysin activity. Neprilysin activity was increased during opioid withdrawal. E-G) Morphine dependence does not increase neprilysin control of nociceptin inhibition of the BLA-Im synapse. E) Example eEPSCs from single representative experiments. F) Bar chart showing percent inhibition from baseline in control and dependent neurons. G) Bar chart showing that the proportion of nociception inhibition prevented by neprilysin activity was not increased in morphine dependence. H-J) When protein kinase A was inhibited with 10µM H89, neprilysin activity was unaltered by morphine withdrawal. H) Example eEPSCs from single representative experiments. I) Bar chart showing percent inhibition from baseline in control and withdrawn neurons. J) Bar chart showing that the proportion of nociception inhibition prevented by neprilysin activity was no different between treatment groups with H89 present. K-M) Activation of adenylyl cyclase with 10µM forskolin in naïve animals enhances neprilysin control of met-enk inhibition. K) Example eEPSCs from single representative experiments. L) Bar chart showing percent inhibition from baseline. Thiorphan boosted the ability of submaximal met-enk (100nM) to inhibit eEPSCs in cells with forskolin present but not in cells with forskolin absent (F_1,8_= 11.26, p=0.01, 2-way ANOVA). M) Bar chart showing the proportion of met-enk inhibition prevented by neprilysin. Neprilysin activity was increased with forskolin present. Bar charts are mean ± SEM. Circles represent individual neurons, with neuron number shown in bars. Data were analysed using 2-way ANOVAs with Bonferroni’s multiple comparisons and unpaired Student’s t-tests. Comparisons on graphs show results from t-tests. See Table S1 for full statistics and analysis.

Hypertrophy in the adenylyl cyclase/PKA signalling pathway is a neuroadaptation revealed in many opioid-sensitive cells during opioid withdrawal (Maldonado et al., 1995, Punch et al., 1997, Williams et al., 2001, Nestler, 2004). Thus, we hypothesised that the withdrawal-induced increase in neprilysin control of peptide activity may result from elevated adenylyl cyclase/PKA signalling. Therefore, we inhibited PKA by incubating morphine-withdrawn slices in H89 (10μM) for 40 mins prior to recording and found that, while neprilysin inhibition still potentiated met-enk inhibition of eEPSCs (Figure 3H-I), the amount of the met-enk response prevented by neprilysin activity was no different between groups (Figure 3J). This suggests that the upregulation of neprilysin activity revealed during opioid withdrawal is produced as a result of increased PKA activity. Thus, increasing adenylyl cyclase activity in neurons from untreated rats should mimic the effect of opioid withdrawal. To test this further, we activated adenylyl cyclase with forskolin (10μM) in slices from naïve animals and found that forskolin potentiated baseline eEPSC amplitudes (68 ± 11.61% increase from baseline, p = 0.02, paired Student’s t-test, n = 5). In the presence of forskolin, submaximal met-enk (100nM) inhibited eEPSCs (Figure 3K-L) and the met-enk inhibition was further increased by neprilysin inhibition in the presence of forskolin (Figure 3K-L). Further, a significantly larger proportion of the met-enk response was prevented by neprilysin activity in neurons with forskolin present (Figure 3I). Forskolin rapidly increases neprilysin breakdown of peptides of within 5-10 minutes of application. The rapid actions of forskolin are consistent with the rapidly developing and then stable withdrawal-induced increase in neprilysin activity that we observed. In control (n=19) and withdrawn cells (n=17), the size of the neprilysin upregulation remained consistent from 2-11 hrs in the morphine-free bath (Control cells: r_17_=-0.3945, p=0.09; Withdrawn cells: r_15_=0.3305, p=0.18, data analysed by two-tailed Pearson’s correlation coefficient). Together, these data indicate that inhibition of PKA prevents the withdrawal induced increase in neprilysin activity and that stimulation of the adenylyl cyclase pathway induces an upregulation of neprilysin activity analogous to the increase observed in withdrawn slices. Therefore, it is likely that enhanced adenylyl cyclase and PKA activity during withdrawal boosts neprilysin activity and, through this, limits peptide regulation of neural circuits in the amygdala.

### Balance is shifted towards inhibition at synapses between Im ITCs and NAc-projecting BLA principal neurons during withdrawal

Having established that peptide regulation of the BLA-Im synapse is turned down during opioid withdrawal due to increased neprilysin activity, we wondered whether peptide signalling is also disrupted at GABAergic projections from Im ITCs onto BLA principal neurons. Given the importance of endogenous opioid signalling in the BLA, this could influence distinct learning mechanisms, such as reward re-evaluation, that occurs in the BLA during opioid withdrawal. We were particularly interested in a subpopulation of BLA principal neurons involved in encoding reward values and mediating reward-seeking behaviours (Stuber et al., 2011, Britt et al., 2012, Namburi et al., 2015, Beyeler et al., 2016, Ramirez et al., 2015). These neurons are characterised by their projections to the NAc. Indeed, studies that optically stimulate, inhibit (Stuber et al., 2011), or lesion (Shiflett and Balleine, 2010) the neural pathway from the BLA to the NAc have confirmed that the activity of NAc-projecting BLA principal neurons is required for animals to correctly process the subjective values of rewards and use these values to guide reward-seeking behaviours. While the Im strongly inhibits randomly sampled BLA principal neurons (Gregoriou et al., 2019), it is not known whether Im neurons specifically target NAc-projecting BLA principal neurons. To identify NAc-projecting BLA principal neurons, fluorescent retrograde beads were injected into the NAc (Figure 4A-B). Then, the Im was electrically stimulated and whole cell recordings were made from BLA principal neurons containing fluorescent beads (Figure 4C-F). BLA principal neurons were characterised by their marked pyramidal morphology with clear differentiation of a thicker apical dendrite from thinner basal dendrites. Synaptic jitter, or the variability of eIPSC latency (the time interval between the end of the stimulation artifact and the onset of the IPSC), indicates whether a synaptic connection is mono- or poly-synaptic, with jitters < 0.7ms signifying monosynaptic connections, and jitters > 0.7ms representing polysynaptic connections (Doyle and Andresen, 2001). The synaptic jitter of each Im-BLA (NAc-projector) synapse was calculated and only neurons with direct connections were kept for further analysis (Figure 4F). Fifty-four NAc-projecting BLA principal neurons were recorded from and of these 49 neurons (91%) had direct inhibitory synaptic connections from the Im (synaptic jitter 0.35ms ± 0.02ms). These data suggest that Im neurons do indeed target the NAc-projecting subpopulation of BLA principal neurons and that withdrawal-induced adaptations could directly affect BLA-NAc reward processes. To address whether peptide control of the Im-BLA (NAc-projector) synapse is also disrupted during opioid withdrawal we tested whether neprilysin more strongly controls nociception inhibition of this synapse in morphine withdrawn vs. control neurons (Figure 4G). Nociceptin was used as we know its actions are limited by neprilysin (Gregoriou et al., 2020) and using opioids was inappropriate as Im terminals are not inhibited by DOR agonism (Winters et al., 2017). Also, MOR activity is likely disrupted during opioid withdrawal, as we saw at the BLA-Im synapse (Figure 1). In both morphine-withdrawn and control neurons, submaximal nociceptin (100nM) inhibited eIPSC amplitude at the Im-BLA (NAc-projector) synapse and this inhibition was potentiated by thiorphan (10µM) (Figure 4H-I). However, a larger proportion of the nociceptin response was prevented by neprilysin in spontaneously withdrawn cells than control (Figure 4J). This indicates that, as we saw at BLA-Im synapses, neprilysin degradation of peptides is enhanced in opioid withdrawn slices at Im-BLA synapses. This disruption of peptide signalling in the BLA, and thus, disruption of GABAergic inhibition of NAc-projecting BLA principal neurons, may disrupt the reward learning processes and resulting behaviours that are mediated by this pathway.

**Figure 4.**
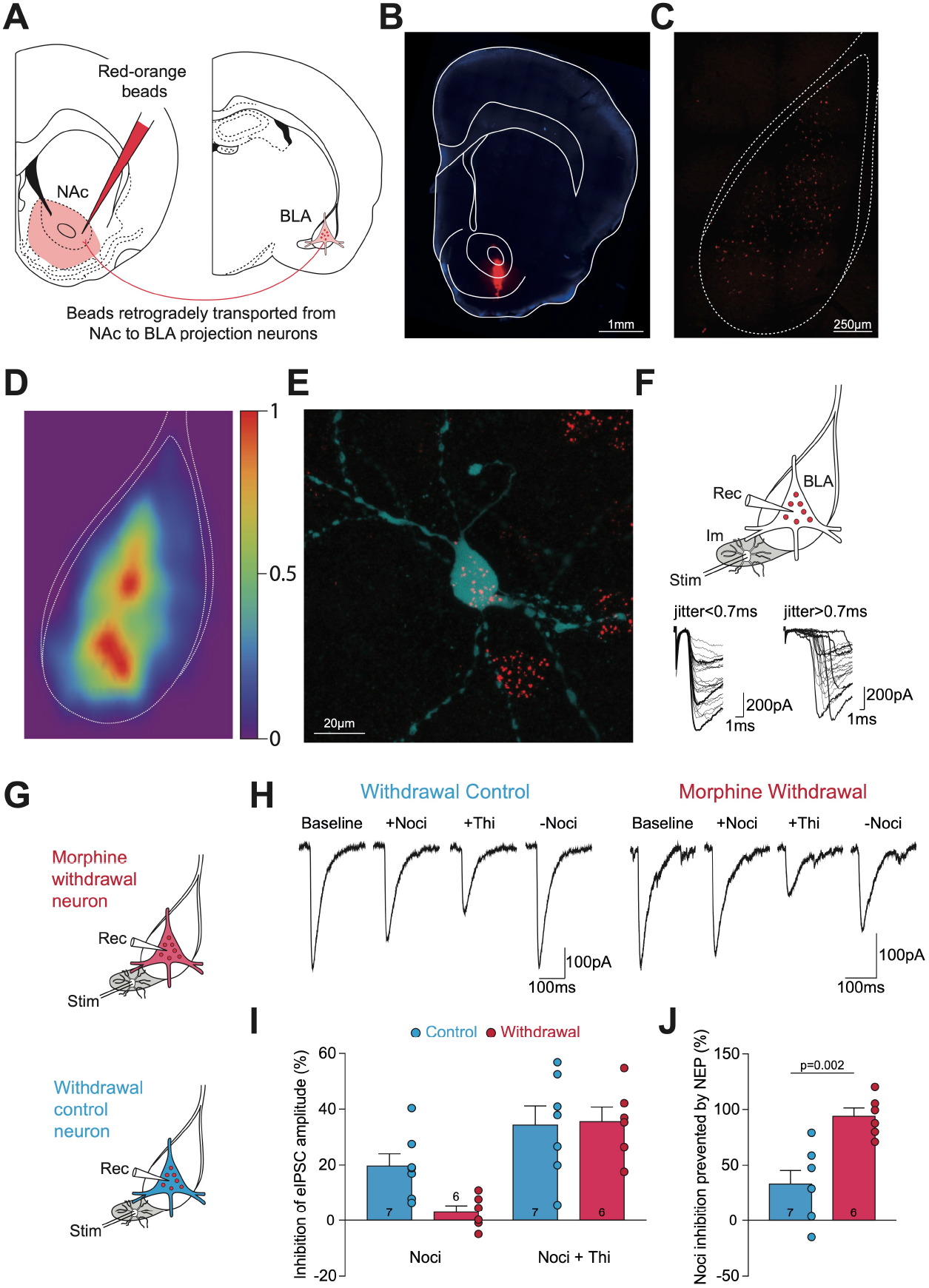
Withdrawal increases GABAergic inhibition of NAc-projecting BLA principal neurons. A) Approach for recording NAc-projecting BLA principal neurons. Schematic shows injection site for fluorescent beads in the NAc and retrogradely transported beads in NAc-projecting BLA principal neurons. B) Representative fluorescence image showing a fluorescent bead injection site in the NAc (red) with DAPI labelling (blue). C) Representative fluorescence image of the BLA showing retrogradely transported beads (red) visible in NAc-projecting BLA principal neurons. D) Heatmap of bead density in intermediate BLA slices. Bregma -1.80, n = 6 slices. In each slice, the position of every NAc-projecting principal neuron was quantified by their X and Y coordinates and then normalised to the top of the BLA. The normalised bead plots were superimposed to generate a heatmap of bead density, with density represented by colour – from purple (low density) to red (high density). E) Biocytin labelled BLA pyramid (blue) positive for retrogradely transported beads (red). F) Typical stimulating site in the Im and recording site in the BLA. Example traces from BLA neurons demonstrating a monosynaptic connection with jitter<0.7ms and polysynaptic connection with jitter>0.7ms. G) Experimental setup for investigation into neprilysin activity at Im synapses onto NAc-projecting BLA principal neurons. H-J) Morphine withdrawal enhances neprilysin control of nociceptin inhibition of the Im-BLA (NAc-projecting) principal neuron synapse. H) Example eEPSCs from single representative experiments. I) Bar chart showing percent inhibition from baseline. I) Thiorphan significantly increased percent inhibition by submaximal nociceptin (100nM) in both control and withdrawn neurons (F_1,11_=50.48, p<0.0001, 2-way ANOVA). While there was no overall effect of treatment group on percent inhibition by submaximal nociceptin, Bonferroni’s multiple comparisons showed there was a trend towards withdrawal significantly reducing the ability of submaximal nociceptin to inhibit eEPSC amplitude (p=0.06). J) Bar chart showing the proportion of nociception inhibition prevented by neprilysin activity. Neprilysin activity was increased by opioid withdrawal. Bar charts are mean ± SEM. Bar charts are mean ± SEM. Circles represent individual neurons, with neuron number shown in bars. Data were analysed using 2-way ANOVAs with Bonferroni’s multiple comparisons and unpaired Student’s t-tests. Comparison on graph shows results from t-tests. See Table S1 for full statistics and analysis.

## Discussion

Here, we reveal a loss of control of synaptic modulation by endogenous peptides due to abnormally high peptidase activity in the Im during withdrawal from chronic morphine treatment. Consequently, inhibition of glutamate release from BLA terminals in the Im is reduced and GABA release from Im terminals onto BLA principal neurons is enhanced during opioid withdrawal (Figure 5). These data established that there are functional increases in neprilysin activity as an adaptive response to chronic drug administration and further, that the increased neprilysin activity relies on withdrawal driven increases in PKA activity. This upregulation of neprilysin activity altered peptide regulation of synapses at multiple sites within the amygdala and for multiple peptides.

**Figure 5.**
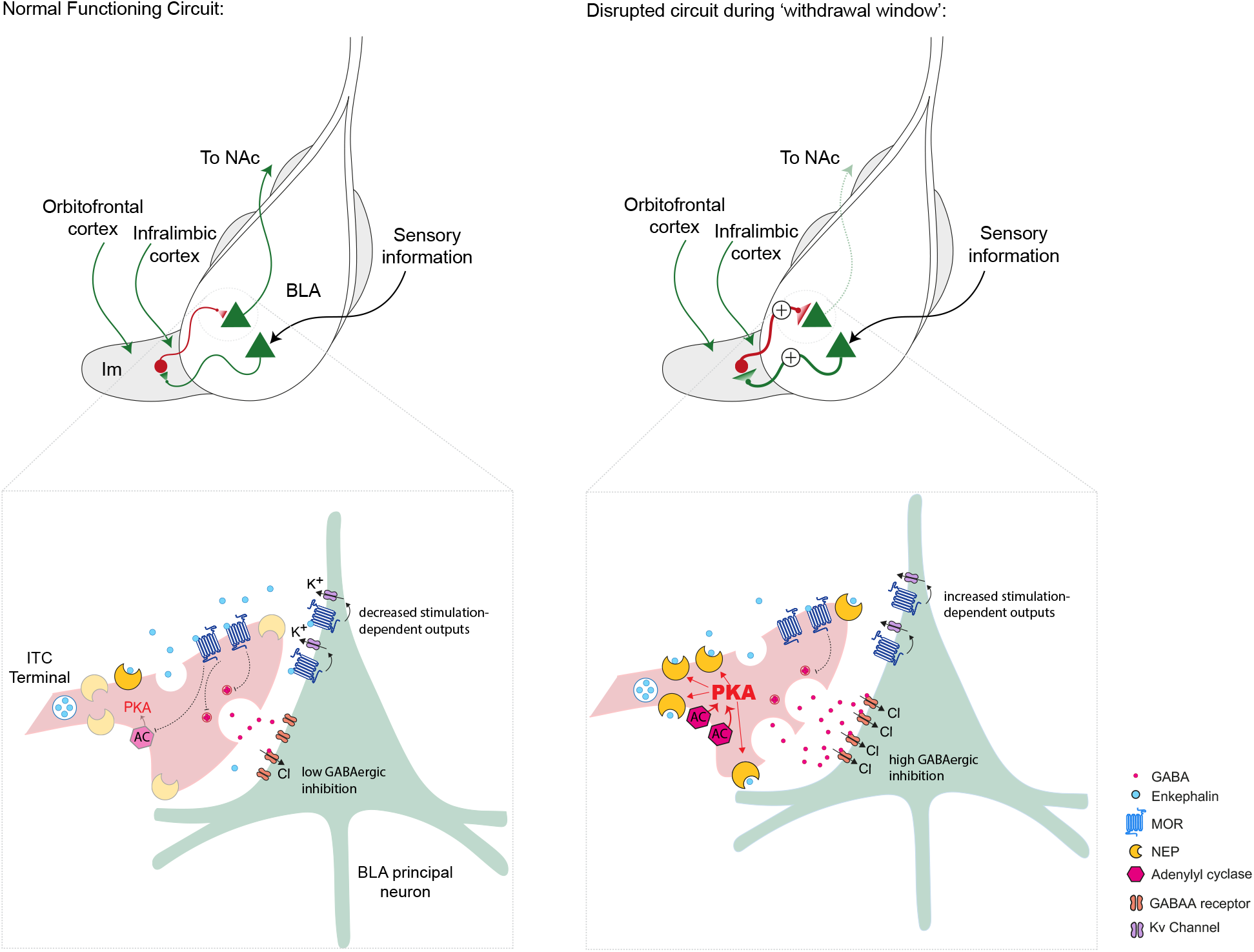
Proposed opioid withdrawal-induced changes to synaptic transmission within the amygdala. Schematic shows withdrawal-induced changes at the circuit level. Under normal conditions, the activation of BLA principal neurons by sensory and also direct cortical inputs generates Im neuronal activity, which drives feedback inhibition onto Im targets, including NAc-projecting BLA reward neurons. During the withdrawal window, reduced opioid inhibition of BLA principal neuron inputs to the Im will increase Im activation. This increased activation and reduced opioid inhibition of GABA release from Im neurons will result in greater feedback inhibition of BLA principal neurons and thus will likely reduce amygdala outputs (i.e., inhibition of BLA outputs to the NAc) and any associated behaviours. Magnification of an Im ITC-BLA(NAc-projecting) principal neuron synapse shows synaptic level changes. Under normal physiological conditions, appropriate stimulation will induce the release of endogenous enkephalins from ITC terminals at concentrations sufficient to overcome peptidase degradation and signal at their presynaptic receptors, inhibiting GABA release from Im terminals, and at their post-synaptic receptors, reducing input-driven activation of BLA principal neurons. During chronic treatment with opioids there is upregulation of adenylyl cyclase/PKA signalling. Withdrawal removes opioid inhibition of the adenylyl cyclase/PKA system and the resulting superactivation rapidly boosts neprilysin peptidase activity. The increased neprilysin activity reduces synaptic concentrations of endogenous enkephalins and impairs the ability of these peptides to signal at their receptors. This increases GABA release from Im terminals and reduces opioid activation of Kv channels on BLA principal neurons.

The capacity for breakdown of opioid peptides is increased in multiple brain regions during opioid withdrawal (Malfroy et al., 1978, Zhou et al., 2001). Here, we used a functional measure of peptidase activity to demonstrate that peptide breakdown is also disrupted in the amygdala and crucially, we showed that this has functional consequences for amygdala neural circuits. To allow us to distinguish withdrawal-induced changes in peptidase activity from changes in receptor function and/or peptide release we used two approaches. Firstly, we studied peptide activation of DOR or nociception receptors, whose activity was not altered by chronic treatment or opioid withdrawal. Secondly, we studied peptidase regulation of exogenous peptide which was not influenced by any changes to endogenous peptide release. This approach allowed us to firmly conclude that opioid withdrawal increased neprilysin activity and thus peptide breakdown, although there may also be other changes, such as reduced responses at the MOR and the possibility of changes in peptide release may occur.

Withdrawal from chronic opioids increases adenylyl cyclase and resultant PKA activity in MOR-sensitive neurons in brain regions such as the periaqueductal gray, striatum and ventral tegmental area (Bonci and Williams, 1997, Nestler, 2004, Williams et al., 2001). This elevation of adenylyl cyclase and PKA is responsible for a range of changes in neuronal activity, including intrinsic excitability, generation of extracellular adenosine and neurotransmitter release (Williams et al., 2001, Bagley et al., 2011). In these experiments, we have discovered that superactivation of this pathway in the amygdala increases the activity of the peptidase neprilysin. How adenylyl cyclase/PKA activity alters neprilysin activity in neurons has not been defined. However, in endothelial cells activation of adenylyl cyclase increases both neprilysin activity and total neprilysin protein levels (Graf et al., 1995), effects that are mimicked by activation of cAMP and PKA (Graf et al., 1995). As we found that opioid withdrawal and activation of adenylyl cyclase with forskolin rapidly increases neprilysin activity, this suggests that PKA phosphorylation of neprilysin, or another regulatory protein, increases the activity of existing neprilysin rather than relying on expression of neprilysin molecules, which takes at least 90 minutes (Stewart and Kenny, 1984, Lemay et al., 1990). Although, additional boosts to neprilysin activity during later phases of withdrawal via increased expression cannot be discounted. The reliance of neprilysin activity on adenylyl cyclase/PKA has several important implications. Firstly, that neprilysin activity could be increased in MOR sensitive neurons in other brain regions where adenylyl cyclase/PKA activity is increased during withdrawal (Nestler, 2004). Consistent with this, elevations of adenylyl cyclase/PKA activity occur in the periaqueductal gray, striatum and ventral tegmental area (Zhou et al., 2001), where there are also elevations in biochemical measures of peptidase activity (Malfroy et al., 1978). The consequence of elevating neprilysin activity will depend on enkephalin function in each brain region. For example, in the striatum it could alter reward learning and behaviours, including state dependent reward consumption, which has recently been shown to rely on enkephalins (Castro et al., 2021). In contrast, in the periaqueductal gray, inhibition of neprilysin significantly attenuates the naloxone-precipitated withdrawal syndrome in rats (Haffmans and Dzoljic, 1987, Maldonado et al., 1992), suggesting that peptidase inhibitors are able to mitigate the reduced peptide function and rescue the impaired peptide signalling that contributes to the withdrawal syndrome. Secondly the PKA dependence suggests that as withdrawal induced increases in adenylyl cyclase/PKA activity persists for at least a week at some synapses (Bonci and Williams, 1996) the adenylyl cyclase/PKA driven neprilysin activity and consequent limitation of peptide levels and actions could be disrupted throughout this time. Finally, that it is possible that other interventions or behavioural states that increase adenylyl cyclase/PKA signalling, such as such as fear learning in the BLA (Goosens et al., 2000, Schafe and LeDoux, 2000, Weeber et al., 2000) or chronic alcohol or cocaine (Nestler, 2004) could enhance neprilysin activity and thus disrupt peptide signalling.

The dialling down of peptide levels in the amygdala during opioid withdrawal will likely limit the extent as well as the temporal and spatial characteristics of peptide regulation at multiple targets, including neurotransmitter release (Winters et al., 2017, Gregoriou et al., 2020) and potassium channel activation (Faber and Sah, 2004, Winters et al., 2017). Given these multiple, and possibly ‘opposing’ targets, the consequences of these changes is likely to be complex, as summarised in Figure 5. However, disruption of the responsiveness/activity and development of plasticity in BLA principal neurons that project to the NAc is likely. In this study we found that Im neurons inhibit nearly all BLA principal neurons that project to the NAc. This suggests that when this cluster of GABAergic neurons is activated by the BLA (Winters et al., 2017), infralimbic cortex (Amir et al., 2011) and, in primates, the orbitofrontal cortex (Ghashghaei and Barbas, 2002), it will inhibit behavioural responses that require activity of BLA-NA pathway, such as cue-guided and reward seeking behaviours (Everitt et al., 1999, Ambroggi et al., 2008, Stuber et al., 2011) (Figure 5). In addition, this GABAergic inhibition could alter the ability of BLA principal neurons to correctly encode, update and/or retrieve reward values (Wassum and Izquierdo, 2015, Malvaez et al., 2019). This is likely because these processes rely upon the ability of BLA principal neurons to form stimulus-outcome associations, which are thought to be mediated, in part, by synaptic plasticity at cortical and thalamic inputs to these cells (Maren, 2005, Bocchio et al., 2017) and is prevented by GABAergic inhibition (Huang and Kandel, 1998, Clem and Huganir, 2010). Given this influence of GABAergic inhibition, it is important that during withdrawal, BLA and cortical activation of the Im will boost GABAergic inhibition onto BLA principal neurons through reduced peptide inhibition of both glutamatergic activation of Im neurons and GABA release from Im neurons (Figure 5). The responsiveness/activity of these BLA neurons may also be increased during withdrawal due to lower opioid activation of voltage gated potassium channels (Faber and Sah, 2004) (Figure 5). Therefore, the disruption of peptide control of the Im may contribute to the impaired ability of animals to correctly evaluate and respond to rewards during both the withdrawal period and beyond, which relies on endogenous opioid actions in the BLA (Wassum et al., 2016). The disruption of peptide function could extend to other peptides that neprilysin degrades, such as dynorphin (Hiranuma et al., 1998), corticotropin releasing factor (Ritchie et al., 2003) and substance P (Matsas et al., 1985), which each play critical and specific roles in opioid addiction (Heinrichs et al., 1995, Gadd et al., 2003, Koob, 2015).

The reduction in peptide function that we observed and thus disruption of amygdala neural pathways could be mitigated using two approaches that we have previously shown enhance endogenous opioid function in the amygdala (Winters et al., 2017). Firstly, inhibition of neprilysin would rescue levels of all neprilysin-sensitive peptides, including enkephalins (Matsas et al., 1984) and nociception (Sakurada et al., 2002), and to a lesser degree other peptides associated with drug use, such as dynorphin (Matsas et al., 1984). This approach is successful in reducing the physical signs of withdrawal when neprilysin inhibitors are injected into the periaqueductal gray (Haffmans and Dzoljic, 1987, Maldonado et al., 1992). The second approach would more selectively mitigate the lower levels of peptides during withdrawal by enhancing receptor responsiveness to their cognate peptides using positive allosteric modulators (PAMs), such as MOR and DOR PAMs (Winters et al., 2017). Both of these approaches may act to close the aberrant learning window that opioid withdrawal opens in the amygdala and should be pursued as viable targets for the development of novel pharmacotherapies for drug addiction.

## Acknowledgments

This work was supported by the National Health and Medical Research Council [Grant APP1047372]

## Author Contributions

Research design: GCG, BLW, EEB

Conducted experiments: GCG, SDP, SP

Performed data analysis: GCG, EEB

Writing of manuscript: GCG, EEB

## Declaration of Interests

There are no sources of conflict of interest

**Figure S1.**
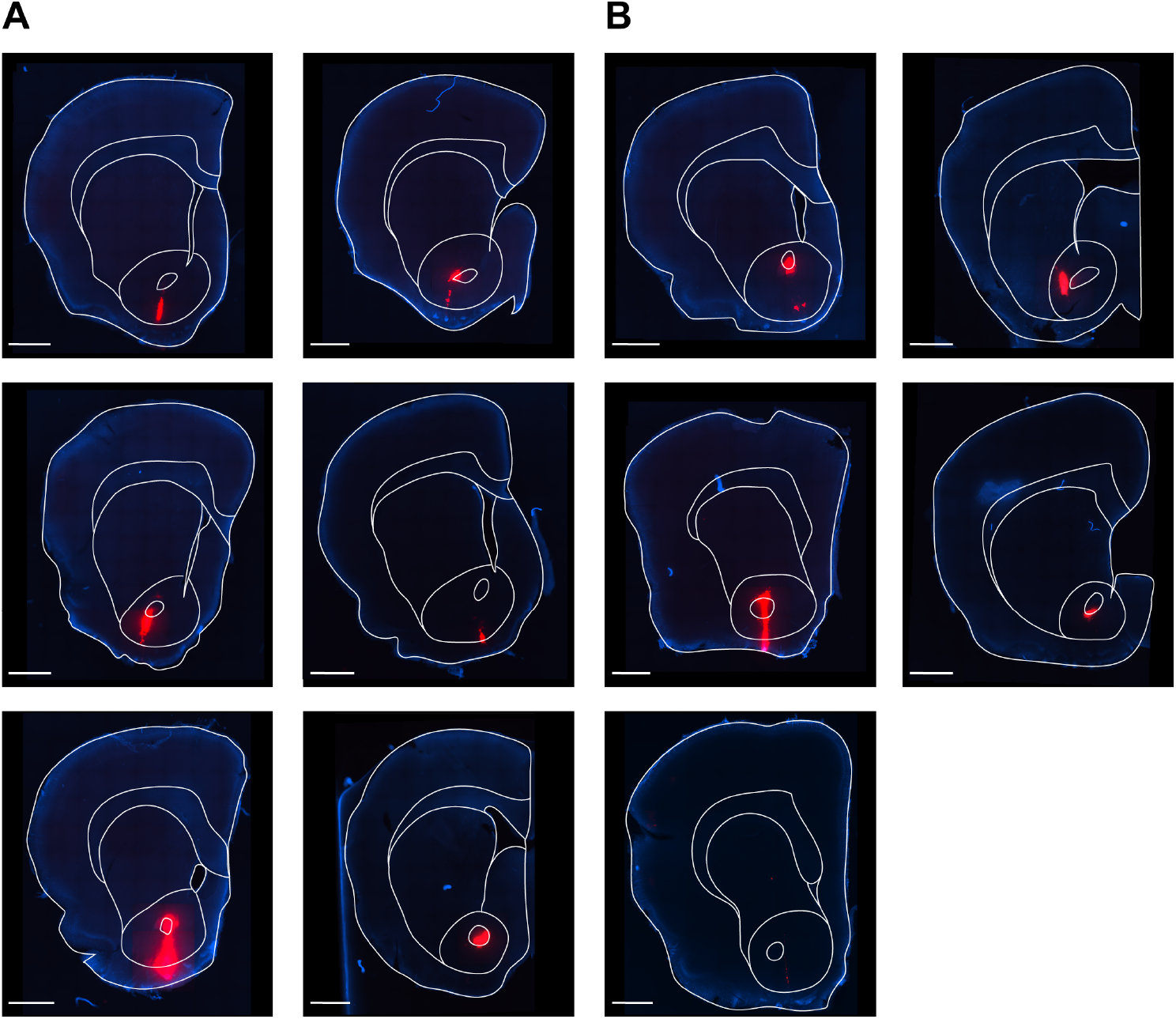
Fluorescent bead injection sites in the nucleus accumbens, related to Figure 4. A-B) Injections of fluorescent red-orange beads were made into the nucleus accumbens. A) Fluorescent images of bead injection sites in nucleus accumbens of vehicle animals (n=6) used in Fig. 4. B) Fluorescent images of bead injection sites in nucleus accumbens of morphine dependent animals (n=5) used in Fig. 4. Fluorescent beads (red) and DAPI labelling (blue). Scale bars, 1mm.

## Online Methods

### General Methods

All experimental procedures were conducted under protocols approved by the University of Sydney Animal Ethics Committee. Experimental procedures were performed on male Sprague-Dawley rats (4–13 weeks old) obtained from the Animal Resources Centre (Perth, Australia). Rats were housed in groups of up to six in a low background noise room. Rats were maintained on a 12/12-hour light-dark cycle with food and water available ad libitum. Room temperature was maintained at 21 (±1) °C.

### Acute brain slice preparation

Coronal slices were prepared according to the methods of (Gregoriou et al., 2020). Rats were anesthetised with isoflurane and killed by decapitation. The brain was rapidly removed from the skull and immersed in ice-cold external solution (in mM: 125 NaCl, 2.5 KCl, 1.25 NaH_2_PO_4_, 2.5 MgCl_2_, 0.5 CaCl_2_, 11 glucose, 25 NaHCO_3_; pH 7.3, 295mOsm/L) bubbled with carbogen (95% O2: 5% CO2).

Coronal brain slices (280μm) containing the rostral amygdala were cut using a VT1200S vibratome (Leica Biosystems, Germany) and then transferred to a holding chamber containing carbogenated cutting solution. In some instances (when animals were over 6 weeks old or when cells were being recorded from the BLA), slices were incubated in a recovery solution (in mM: 93 NMDG, 2.5 KCl, 1.2 NaH_2_PO_4_, 30 NaHCO_3_, 20 HEPES, 25 glucose, 2 thiourea, 3 sodium pyruvate, 5 sodium ascorbate, 10 MgCl_2_, 0.5 CaCl_2_; pH 7.3, 300-310mOsm/L) for 10 mins prior to being transferred to the holding chamber. Occasionally slices had to be pre-treated with drugs before recording. In these cases, drugs were included in the external solution (noted in results where applicable). For all experiments, once submerged in cutting solution, slices were incubated at 34°C for at least 1 hour and then stored at room temperature until used for recording.

### Electrophysiology

For recording, brain slices were transferred to a recording chamber and continuously superfused at 2ml/min with artificial CSF (aCSF) of composition (in mM) 125 NaCl, 2.5 KCl, 1.25 NaH_2_PO_4_, 1 MgCl_2_, 2 CaCl_2_, 11 glucose and 25 NaHCO_3_, saturated with carbogen and heated to 32-34°C. Brain slices were visualised with a BX51WI upright microscope (Olympus, Japan) fitted with Dodt gradient contrast optics (Thorlabs, USA). Neurons of interest were readily identified by their location and morphology. Fluorescent neurons were visualised using an X-cite series 120Q excitation light source (Excelitas Technologies Corp, USA) and appropriate filters. To verify correct targeting of neurons, either the electrophysiological properties of cells were examined, or slices were fixed following recording and kept for post-hoc staining (see below). Whole-cell voltage-clamp recordings were made using patch electrodes (2-5 MΩ) pulled on a P-1000 Micropipette puller (Sutter Instruments, USA).

Whole-cell voltage-clamp recordings were made from neurons clamped at -70 mV using patch pipettes filled with internal solution containing (in mM): 140 CsCl, 5 HEPES, 10 EGTA, 2 CaCl_2_, 2 MgATP, 0.3 NaGTP and 3 QX-314-Cl (pH 7.3, 280-285mOsm/L). 0.1% Biocytin was routinely added to the internal solution. The liquid junction potential of -4mV was not corrected. In all experiments, series resistance (<20MΩ) was compensated by 60% and continuously monitored. Experiments in which series resistance changed by >20% were excluded from analysis.

Electrically evoked synaptic currents were elicited via a bipolar stimulating electrode placed in various positions depending on the synaptic input studied. To record evoked excitatory postsynaptic currents (eEPSCs) at BLA-Im synapses, the stimulating electrode was placed close to the basomedial edge of the BLA and the GABA_A_ receptor antagonists picrotoxin (100μM) and gabazine (10μM) were included in the aCSF to block fast inhibitory synaptic transmission. To record evoked inhibitory postsynaptic currents (eIPSCs) at Im-BLA synapses, the stimulating electrode was placed within the Im and the AMPA receptor antagonist CNQX (10μM) was used to block fast excitatory synaptic transmission.

### Electrophysiology data acquisition and analysis

All recorded electrophysiological signals were amplified, low pass filtered (5 kHz), digitised and acquired (sampled at 10kHz) using Multiclamp 700B amplifier (Molecular Devices, USA) and online/offline analysis was performed with Axograph Acquisition software (Molecular Devices, USA).

For all experiments examining synaptic transmission, synaptic currents were evoked every 15 seconds by delivery of paired stimulating pulses (1-100V, 100μs, 50ms inter-pulse interval), expect for endogenous opioid release experiments where a moderate stimulus protocol was used (1 – 100V, 5 stimuli at 150Hz, followed by 1 test pulse 200ms later). Stimulus intensity was defined for each experiment as the voltage at which a synaptic current could be easily and consistently distinguished yet yielded sub-threshold current amplitudes. All currents were analysed with respect to their peak amplitude. Peak amplitude was quantified as the mean peak amplitude of four-eight consecutive currents, after responses reached a stable plateau. The effects of exogenously applied drugs were indirectly measured via their effects on the peak amplitude of synaptic currents. The peak amplitude of the currents elicited by pulse 1 of the paired stimuli, or the test pulse of the moderate stimuli, were used for analysis of drug effects. Drug effects were quantified as the percentage change in mean peak amplitude between drug superfusion and the average baseline. The average baseline was an average of the peak amplitude of currents at the beginning of each experiment and the peak amplitude of currents at the end of each experiment (upon reversal of drug effects by antagonist or drug washout). Reversal of drug effects was calculated as the percentage of the baseline amplitude recovered by drug washout or antagonist application. In experiments where antagonists increased peak amplitude of currents, drug effects were calculated from the regular baseline. To determine whether synaptic connections were monosynaptic, the synaptic jitter, or the variability of latency to peak onsets, was measured. Latency to peak onset was defined as the time from the end of the stimulation artefact to 1% of the eIPSC peak. Synaptic jitter of < 0.7ms indicates that a connection is monosynaptic (Doyle and Andresen, 2001), thus, only neurons with a jitter < 0.7ms were kept for further analysis. The synaptic jitter of a response was calculated as the standard deviation of the peak onsets.

### Drugs and reagents

Stock solutions of all drugs were diluted to working concentrations in aCSF immediately before use and applied to brain slices by superfusion. The stock solutions for all drugs, except thiorphan, were made in Milli-Q water. Thiorphan was dissolved in DMSO to achieve a working DMSO concentration of 0.01%. Drugs and reagents were obtained from the following sources: CNQX, DAMGO, SR-95531 (gabazine), thiorphan and bestatin were from Abcam (Cambridge, UK). Picrotoxin, methionine-enkephalin (met-enk) and captopril were from Sigma (St. Louis, Missouri, USA). CTAP, [D-Ala2]-Deltorphin II (Delt II), ICI-174864, naloxone and nociceptin were from Tocris (Bristol, UK). H-89 and forskolin were from Cayman Chemicals (Ann Arbor, Michigan, USA) and morphine base was purchased from Sun Pharma (Port Fairy, Victoria, Australia).

### Chronic morphine treatment

Morphine dependence was induced using the chronic morphine treatment regimen described previously (Bagley et al., 2011). Rats received a series of subcutaneous injections of a sustained release emulsion containing 50mg of morphine base (100mg/kg, Sun Pharma, Australia) suspended in a vehicle solution comprised of 0.1ml mannide monooleate (Arlacel A), 0.4 ml light liquid paraffin and 0.5ml 0.9% w/v NaCl. Injections of warmed emulsion were made while animals were under light isoflurane anaesthesia on days 1, 3 and 5. A group of rats were injected with a morphine-free emulsion on the same schedule and used as controls. Vehicle- and chronic-morphine treated rats were used on days 6 and 7 for experiments. The animal used on day 6 alternated between chronic morphine treated and vehicle throughout experiments.

### Withdrawal behaviours

To test for dependence, a group of rats were injected with the opioid receptor antagonist naloxone hydrochloride (5mg/kg, intraperitoneal). Withdrawal signs including jumping, wet-dog shakes, tremor and teeth chattering were counted before and after withdrawal was precipitated.

### Stereotaxic surgery

Rats were placed in induction chamber and administered 5% isoflurane. When deeply anaesthetised, rats’ heads were shaved and placed in stereotaxic apparatus (Model 942, Kopf Instruments, USA) while maintaining incisor bar to achieve flat skull position. Subcutaneous injection of caprofen (5 mg/kg, Cenvet, Australia) and bupivivaine (5%, Cenvet, Australia) were made under the skin of the injection site. A single incision was made down the centre of the rats’ heads to expose the skull. Holes were drilled above the NAc and bilateral injections of fluorescent beads (1:1 dilution in saline, red-orange fluorescent microspheres 565/580, 0.04μm, ThermoFisher Scientific) were made by hydraulic pressure at 10nL/min for 1 min at stereotaxic coordinates: AP: +1.8 mm, ML: 1.4 mm, DV: -7.2 mm (horizontal skull, reference from bregma). Injections were made with glass micropipettes (30 – 50μm tip, Drummond Scientific, USA) pulled with a PC-10 micropipette puller (Narishige, Japan). On completion of the injection, the micropipette was left in place for a further 15 mins before retraction. Following pipette withdrawal, skull openings were sealed with bone wax (Coherent Scientific, Australia) and the incision was closed with silk sutures. Rats were given cephazolin (100mg/ml, Hospira, Cevnet, Australia) and saline post-surgery and allowed to recover for 7 days before used for experiments.

### Immunohistochemistry

Slices were prepared for immunohistochemistry from perfused animals or following electrophysiological experiments for post-hoc staining. For perfusion, animals were deeply anaesthetised with isoflurane and/or a lethal dose of pentobarbital sodium (120–150mg/kg, intraperitoneal). Once reflexes were abolished, animals were perfused through the ascending aorta with 3,000 units per ml heparin in a 0.5% NaNO_2_/0.9% saline solution followed by 4% paraformaldehyde (PFA) solution in 0.1M phosphate-buffered saline (PBS, pH 7.4). Brains were removed and fixed overnight in 4% PFA (4°C). Brains were washed three times with PBS and then stored for no more than 1 week in PBS (4°C) before sectioning. Coronal sections (40μm) containing the amygdala were cut using a vibratome (Leica Biosystems, Germany). Sections were stored in a cryoprotectant solution comprised of 40% PBS, 30% glycerol, 30% ethylene glycol cryoprotectant at −30°C until required for immunohistochemistry. Following electrophysiological experiments, coronal slices (280μM) containing the amygdala were fixed overnight in 4% PFA (4°C). Slices were washed three times with PBS and then stored in cryoprotectant solution at −30°C until required for immunohistochemistry.

Prior to staining, cryoprotectant was removed from slices by washing three times with PBS, then incubated for 30mins at room temperature in 10% donkey serum and 0.5% BSA in PBS to block non-specific binding. Primary antibodies for met-enk (Millipore; Rabbit anti-Met-enkephalin; ab5026, RRID:AB_91644; [1:200]), MOPR (Aves Labs; Chicken anti-MOPR; [1:5000]), and NEP (R&D Systems; Donkey anti-goat; AF1126; lot: JGIO116081, RRID:AB_2144426; [1:80]) were diluted in 0.1% BSA/0.25% Triton X-100 in PBS and sections were incubated overnight (4°C). Secondary antibodies for rabbit (Abcam; Donkey anti-Rabbit Alexa Fluor 568; ab175692; lot: GR322655-2, RRID:AB_2884939; [1:500]), chicken (Jackson ImmunoResearch; Donkey anti-Chicken Alexa Fluor 647, RRID:AB_2340379; [1:500]) and goat (Abcam; Donkey anti-goat Alexa Fluor 488; ab150133; lot: GR312722-2, RRID:AB_2832252; [1:500]) were diluted in 0.1% BSA in PBS, and incubated 2 h at room temperature (light protected). The nuclear stain DAPI (1:500, ThermoFisher Scientific) was added for the final 30 min of this incubation. Sections were then washed three times in PBS and mounted onto slides using ProLong Gold Antifade (Life Technologies).

For post hoc staining of brain slices (280μm) used in electrophysiological experiments and containing cells filled with 0.1% biocytin (Sigma, St. Louis, Missouri, USA) and fluorescent beads (ThermoFisher Scientific, Massachusetts, United States), slices were briefly washed before being incubated for 1 h at room temperature in 10% goat serum, 0.5% BSA and 0.3% Triton X-100 in PBS. Steptavidin conjugated antibody (Alexa Fluor 647 streptavidin, 1:2000, ThermoFisher Scientific S32357) was diluted in 1% BSA/0.1% Triton X-100 in PB and incubated for 2hr at room temperature (light protected). DAPI (1:500, ThermoFisher Scientific) was added for the final 30min of this incubation period. Slices were then washed three to four times (10 min) with PBS and mounted onto slides using Fluoromount-G (SouthernBiotech).

Sections were visualised using a Zeiss LSM800 Meta confocal microscope (lasers 405, 488, 561nm; Carl Zeiss, Germany) and a Leica SP8 STED confocal microscope (lasers 405, 561, 647nm; Leica, Germany). Images were taken sequentially with different lasers using low power dry 10x (numerical aperture (NA) 0.45) and 20x objectives (NA 0.8), as well as high power oil immersion 63x (NA 1.40) objective. Single images were collected using the 10x objective and Z-stacks were collected for the 20x and 63x objectives, respectively

### Data collection and analysis

All data are presented as the mean ± standard error of the mean (SEM). No statistical methods were used to predetermine sample size, however the numbers obtained are similar to those generally used within the field. Numbers vary due to effect size, variability and difficulty of experiment. Data distribution was assumed normal in all cases. Statistical analysis was determined before any results were obtained. All statistical analysis was performed using GraphPad Prism 6. Tests and results are detailed in Table S1. Data were considered significant is p<0.05.

---

**Table S1.**
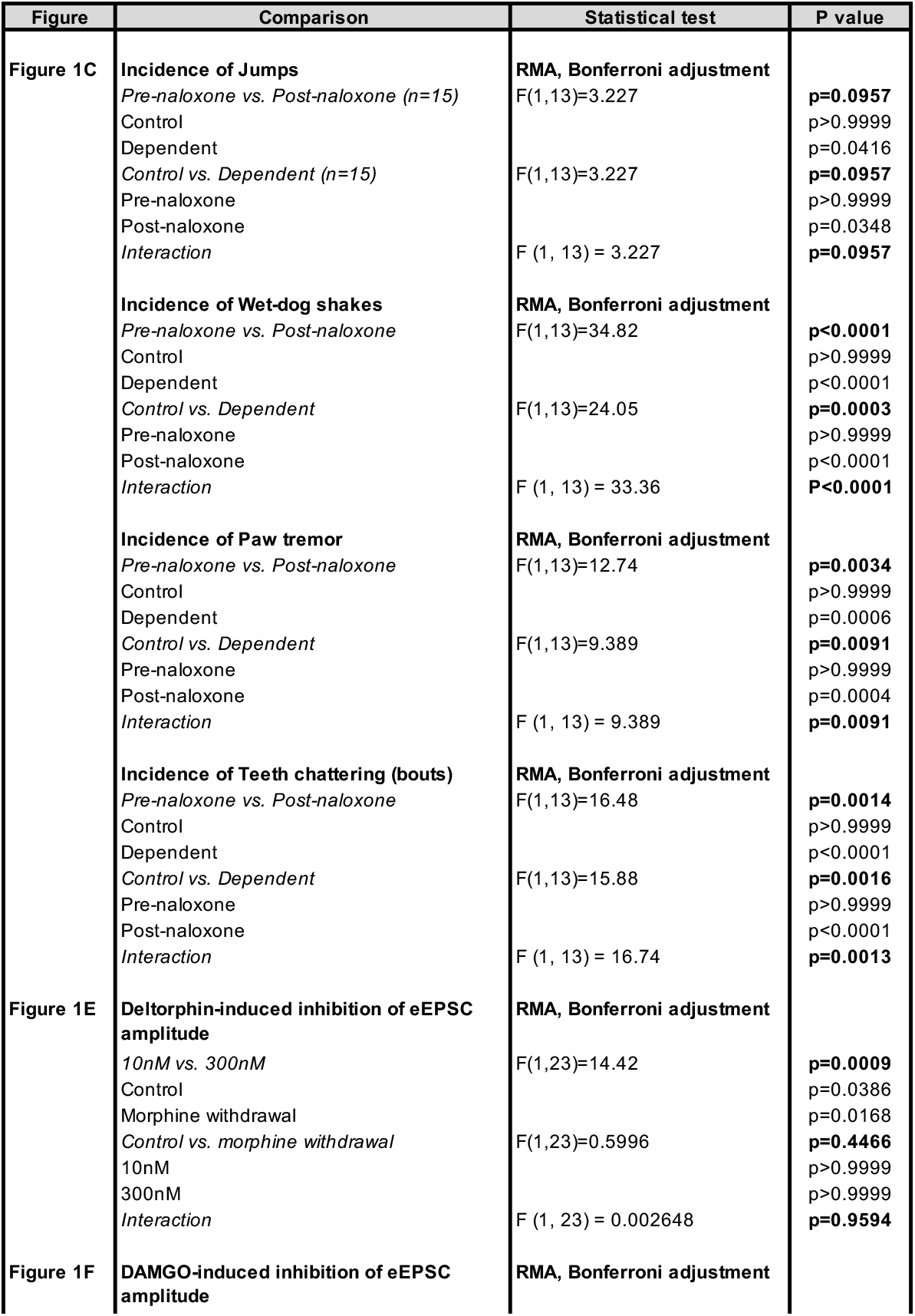
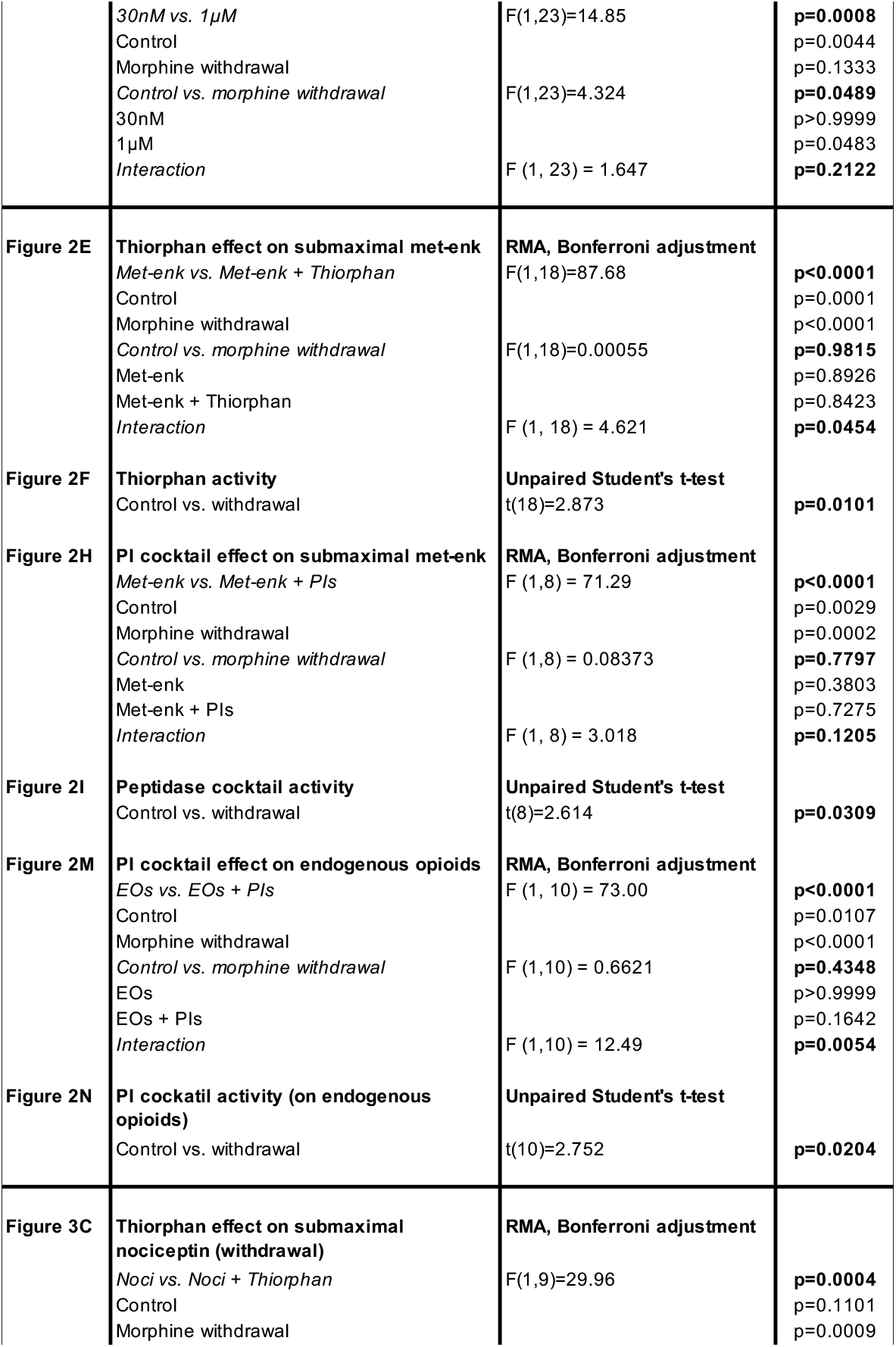
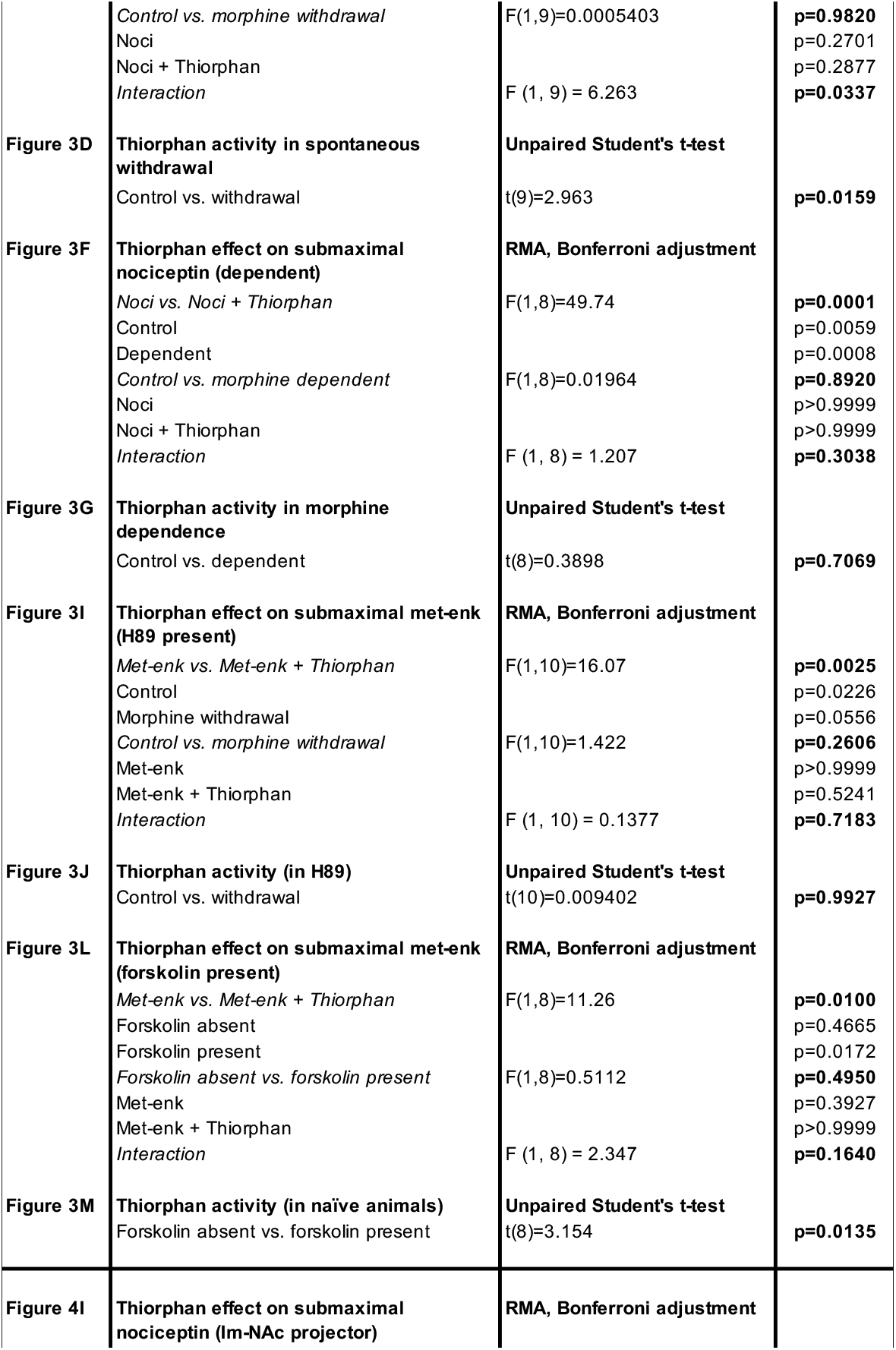
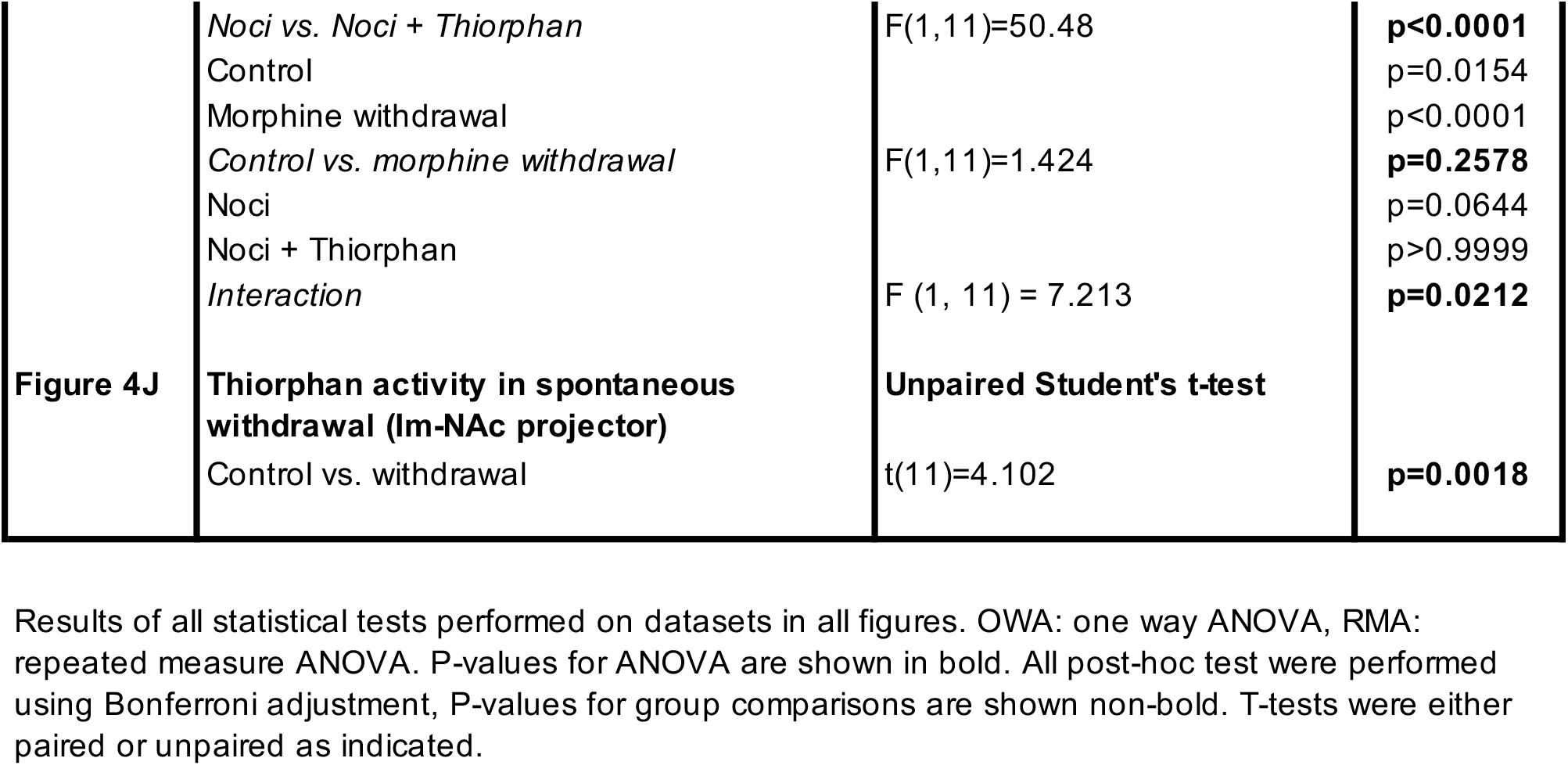
All statistical results.

